# The special and general mechanism of cyanobacterial harmful algal blooms

**DOI:** 10.1101/2021.11.22.469516

**Authors:** Wenduo Cheng, Somin Hwang, Qisen Guo, Leyuan Qian, Weile Liu, Yang Yu, Zhenghao Li, Tianji Wang, Yi Tao, Huansheng Cao

## Abstract

Cyanobacterial harmful algal blooms (CyanoHABs) arise as cyanobacteria dominate phytoplankton communities when nutrient levels increase from oligotrophic state. From a wholistic perspective, this longstanding altered phytoplankton structure results from two conditions: one sufficient condition that cyanobacteria can grow maximally with elevated nutrients; one necessary condition that co-living algae cannot grow fast or dominate at the same levels. The sufficient condition, the ‘special’ mechanism of CyanoHABs at the population level, has been established as the synergistic interaction between superior cyanobacterial ecophysiology and elevated nutrients. But it is unknown how these functions arise or whether they are under directed evolution to water eutrophication. The necessary condition, the ‘general’ mechanism of CyanoHABs at the community level, is little understood: why co-living algae cannot form blooms as cyanobacteria? Literature and bioinformatics analyses show that the superior ecophysiology undergoes no directed positive evolution to worldwide eutrophication in general or any local eutrophic waters in particular; instead, these functions are under strong purifying selection and likely acquired through early adaptive radiation in nutrient-deficient conditions, as functions enabling extant cyanobacteria to occupy other niches. The general mechanism turns out to be quite straightforward: cyanobacteria are simple life forms and thus have lower per capita nutrient demand for growth, compared to co-existing eukaryotic algae in cell size and structure, genome size, size of genome-scale metabolic networks, cell content, nutrient requirement. Lower nutrient demand is proved by existing field nutrient supplementation. Both the special and general mechanisms of CyanoHABs are tentative frameworks awaiting further theoretic improvement and empirical assessment.

## I. Introduction

CyanoHABs are one of the most profound environmental hazards in modern human history in terms of their global geographical scale (Paerl et al. 2020), longstanding duration (over a century) (Francis 1878, Lathrop and Carpenter 1992), and tremendous economic loss (Steffensen 2008). The mere fact that they are still intensifying and expanding under the global climate change (Chapra et al. 2017, Gobler 2020, Paerl et al. 2020) attests the fact CyanoHABs are still not fully understood, such that their ecology remains ‘complicated and confusing’ (Wilhelm et al. 2020). To demystify CyanoHABs, one needs to look at them from a wholistic perspective, considering all the main factors: bloom-forming cyanobacteria, other co-existing algal species, and eutrophic conditions. So far, it has rarely been seen blooms of other groups of algae appear in the same eutrophic waters as cyanobacteria (e.g., Heisler et al. 2008, Steffen et al. 2014, Cook et al. 2020). Considering this upscaled context, the main question now is why cyanobacteria, other than other algae, form blooms in the same eutrophic waters. Apparently, this overarching question can be broken down to two, each of which constitute the sufficient and necessary conditions: (1) how cyanobacteria form blooms in the eutrophic waters, and (2) why co-existing algae do not form blooms in the same eutrophic waters? The second question is equivalent to whether other algae can form blooms under the same eutrophic conditions, whether there are enough supplies of nutrients for them to bloom, or both.

To address these questions, we formulate a perspective primarily based on the comparatively advantage of cyanobacteria in nutrient demand. Simply put, it postulates that cells of cyanobacteria have simple cell structures and small sizes and therefore require less input of energy and matter to sustain and grow than those of other species, and as a result, can grow fast when nutrient level increases in the eutrophic waters. This simple life structure and operation in turn results from adaptive evolution during nutritiously prohibiting environment of early Earth. This perspective will be established by addressing the sufficient and then the necessary condition, respectively.

## II: The special mechanism of CyanoHABs

From an energy and matter perspective, the ubiquitous presence of CyanoHABs attests the fact that bloom-forming cyanobacteria have metabolic machinery able to effectively assimilate external nutrients and convert them to biomass, aided by favorable weather or climate conditions (O’Neil et al. 2012, Zhang et al. 2012). That is to say that CyanoHABs are the outcomes of the synergistic interactions between superior ecophysiology of cyanobacteria and eutrophic conditions. Indeed, both factors have been well established by field work, laboratory simulations, genomics and metatranscriptomics. Specifically, the roles of nutrients, either trace elements or macronutrients, have been recognized as important drivers for CyanoHABs in a consensus and large survey (Heisler et al. 2008, Beaulieu et al. 2013). Meanwhile, the metabolic pathways importing and converting available nutrients have been found to be more complete and have more copies of genes encoded in the genomes of bloom-forming cyanobacteria than non-blooming cyanobacteria (Cao et al. 2020a, Du et al. 2021) (Fig. 1). These metabolic pathways and their genomes are now curated in a webserver CyanoPATH (Du et al. 2021). Most of the functions are peripheral and involved in nutrient uptake and precursor processing to feed central metabolism, and the rest are in the central metabolism converting metabolites for synthesis of macromolecules in anabolism. Interestingly and expectedly, the gene expression levels of these pathways are correlated with their completeness (Cao et al. 2020a). To sum up, at the population level, the synergistic interactions between superior metabolic capabilities and elevated nutrients suffice as a working mechanism of CyanoHABs.

**Figure 1.**
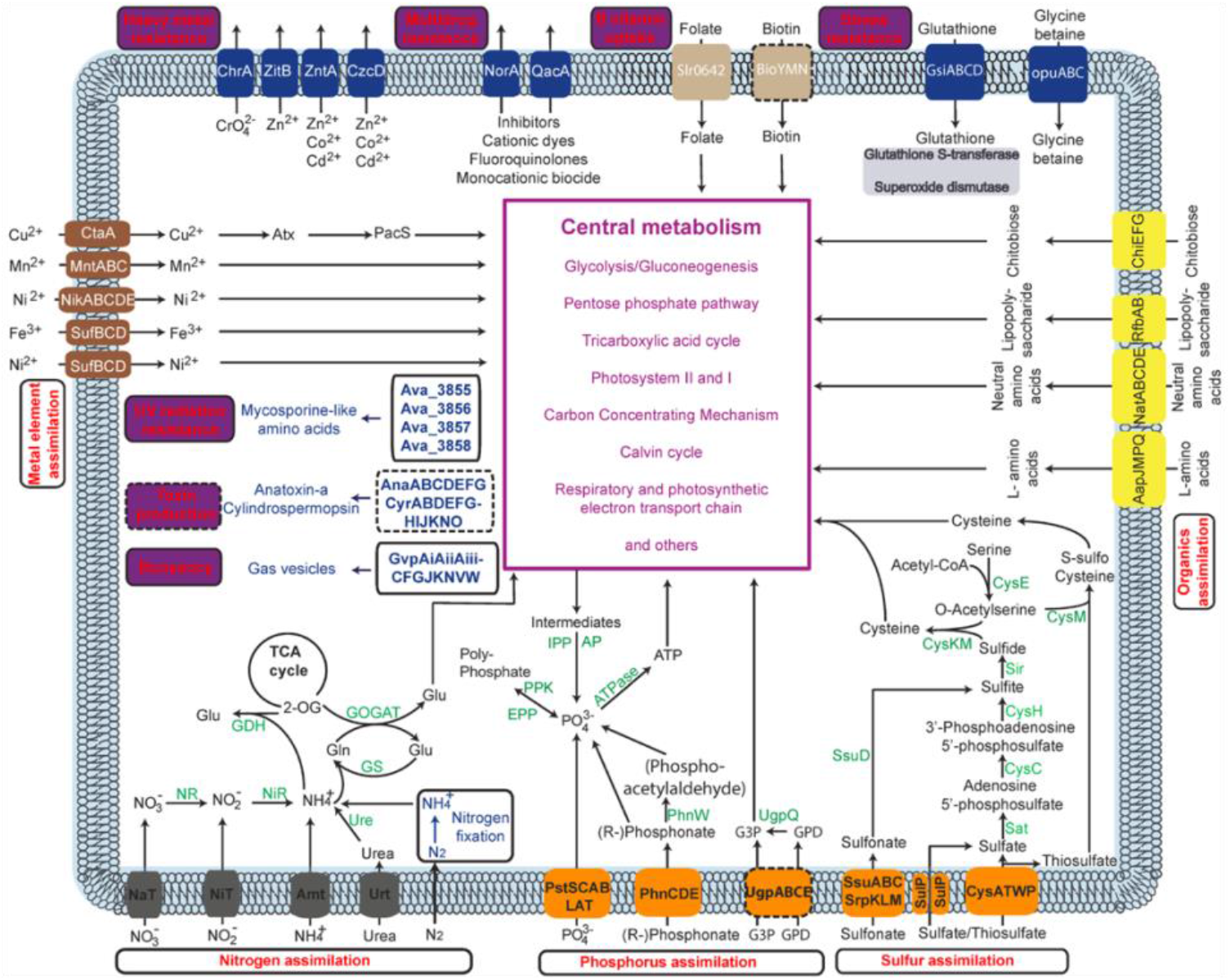
The core and query pathways in *Aphanizomenon flos-aquae* NIES-81. The pathways labeled with dashed borders are not complete (due to the absence of required components) in this strain but may be complete in others.

## III: Is there *a priori* directed evolution toward water eutrophication?

Given their global persistence for over a century (Francis 1878, Lathrop and Carpenter 1992), one may wonder whether there is prior adaptive evolution in cyanobacteria toward water eutrophication to form bloom, or CyanoHABs are simply ecological consequences of synergistic interactions between superior pre-equipped biological functions and elevated nutrients. We think the latter is more likely than the former. First, short-time whole-lake experiments in fertilization or nutrient reduction (Schindler et al. 2016, Wilkinson et al. 2018, Molot et al. 2021) have indisputably shown the causality of eutrophication for CyanoHABs on an ecological time scale, allowing no time for evolution. Second, all evolution of microbes is passive, rather than proactive, response to environmental or genetic changes which are unforeseeable to them, and therefore they cannot proactively prepare themselves through directed evolution.

Third, we confirmed there is no directed evolution toward CynaoHABs, by calculating the metric, dN/dS (ratio of non-synonymous to synonymous mutations), for all homologous genes in 50 blooming species of *M. aeruginosa* genomes from different countries. Indeed, almost all homologous genes display dN/dS < 0.8, with a median of 0.32 (Fig. 2A), indicating they are all under negative (purifying) selection not positive (divergent) evolution. When grouped into the functional pathways curated in CyanoPAHT (Cao et al. 2020b), the homologous genes associated with CyanoHABs displayed the same levels of purifying selection pressure as those in core metabolism and housekeeping processes (e.g., translation and transcriptional regulation), with a mean of dN/dS at 0.32 (Fig. 2B). This strong purifying selection suggests these gene functions not only are important functions such that cells cannot afford them to be comprised by mutations. Furthermore, the phylogenetic analysis shows that the *M. aeruginosa* strains from different countries tend to align in the same clusters (Fig. 2C). This complex phylogeny clearly shows among other things there is no adaptation to local water conditions. In other words, all bloom-forming cyanobacteria share the set of metabolic capabilities which enable them to grow well and dominate eutrophic waters irrespective of local peculiarity in water conditions.

**Figure 2.**
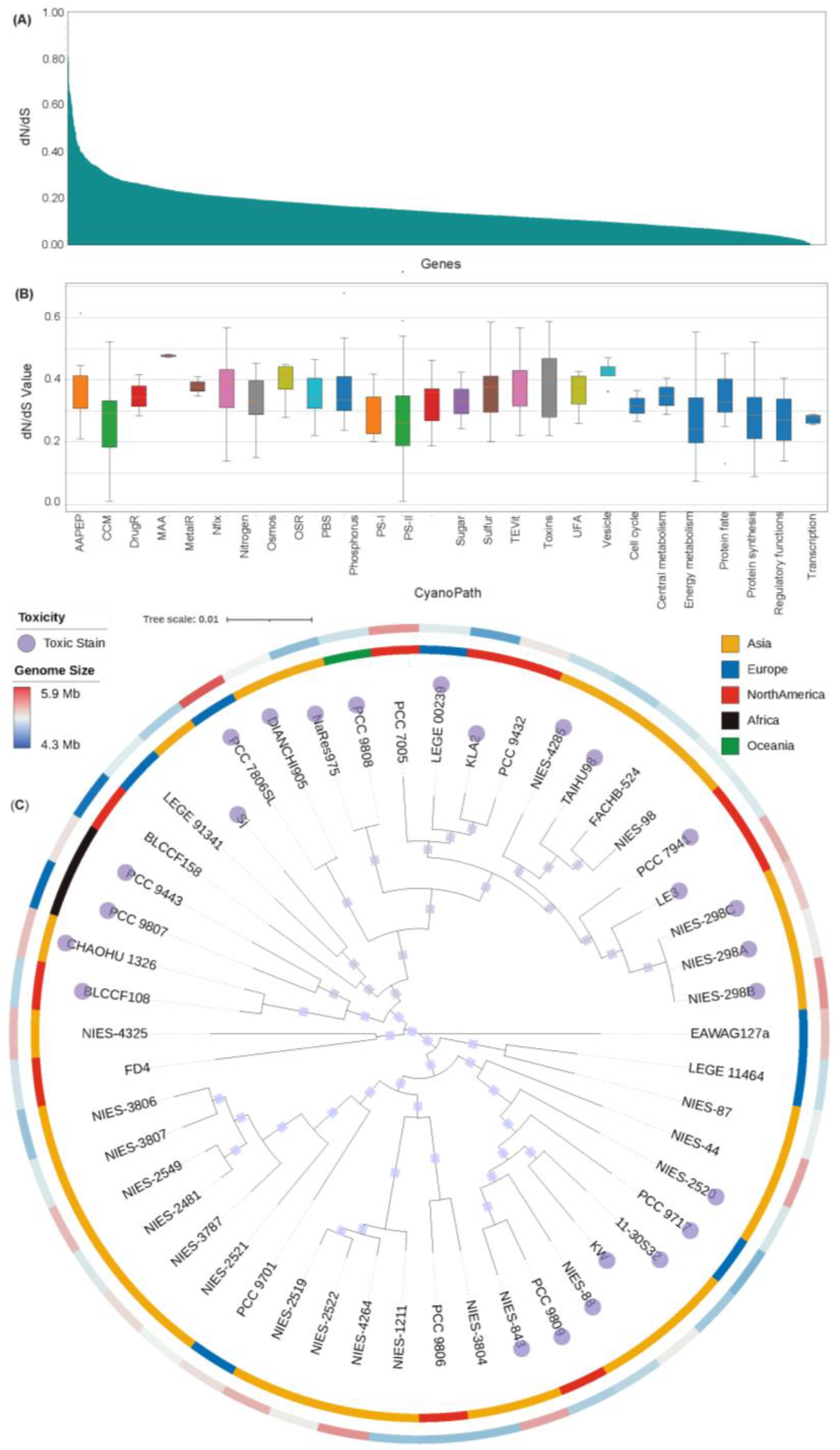
dN/dS for all homologous genes in 50 genomes of *M. aeruginosa*. (A) An overall distribution of dN/dS for all homologous genes; (B) the dN/dS for genes in different groups of functions in CyanoPATH. dN/dS values were calculated by CODEML from the PAML 4.9j package with default settings (Yang 2007). AAPEP: uptake of amino acids and peptides; CCM: CO_2_-concentrating mechanism; DrugR: antibiotics resistance; MAA: UV radiation; MetalR: heavy metal resistance; Nfix: nitrogen fixation; Nitrogen: nitrogen utilization; Osmos: osmos homeostasis; OSR: oxidative stress resistance; PBS: phycobilisome; Phosphorus: inorganic/organic utilization; PS-I/PS-II: photosystem I and photosystem II; Sugar: sugar assimilation; TEVit: assimilation of trace metals and vitamins; Toxins: cyanotoxins; UFA: unsaturated fatty acids; Vesicle: gas vesicles. The last seven columns colored dark blue represent essential gene sets (Rubin Benjamin et al. 2015). (C) The phylogeny of the 50 *M. aeruginosa* strains collected from different countries.

Last, there are not enough raw materials, mutations, for adaptive evolution to be accumulated during one annual cycle of blooms. The spontaneous mutation rate (*µ*) in cyanobacteria is estimated at 10^−7^ per genome per generation (García-Villada et al. 2004, López-Rodas et al. 2007, Osburne et al. 2011, Segawa et al. 2018). For a lake of a million m^3^ (i.e., with a surface area of 1 million m^2^ with a depth of 1 m where cyanobacteria are mostly concentrated), with initial cyanobacteria 20 cells/L growing to 10^8^ cells/L after 25 doublings (*t*) at a growth rate 0.05-1.1 d^-1^ in an annual cycle (bottlenecked in fall and winter), this gives an effective population size (*N_e_*) of 10^8^ cells, based on the equation 15 in Wilson et al. (2006). So, the number of beneficial mutations to be fixed can be approximated by, assuming a beneficial rate (*s*) at 0.02 and a rate of fixation (2*s*) at 0.04, *N_e_ µ t* 2*s* (10^8^ × 10^−7^ × 25 × 2× 0.02 = 10). Therefore, for a lake with a surface area smaller than 0.1 million m^2^, each year there is less than one beneficial mutant will occur, which makes prior adaptive evolution highly unlikely. But large lakes are likely to accumulate beneficial mutations and better-growing mutants.

## IV. Superior functions are shaped during long adaptive radiation on early Earth

The global ubiquity and expansion of CyanoHABs indicate the superiority of metabolic capabilities. The fact these superior functions are under negative selection, instead of assumed positive selection, suggests the metabolic capabilities were well established before the era of water eutrophication. This brings another critical question: how cyanobacteria acquired these functions? Revisiting the origin and evolution of cyanobacteria on early Earth provides important clues: these superior functions are shaped during their course of adaptive radiation.

Originating in an anoxic biosphere of early Earth, cyanobacteria are likely to evolve in scarcity of macronutrients such as phosphorus and nitrogen (Falkowski 1997, Bjerrum and Canfield 2002, Kipp and Stüeken 2017, Reinhard et al. 2017) and trace metals (Anbar and Knoll 2002, Saito et al. 2003). This humble beginning of life leads to primitive metabolic networks (Goldford and Segrè 2018), which suggests that cyanobacteria could only obtain minimal energy and matter in both variety and amount to support cell survival and division (Schopf 2002). Arguably, their nutrient requirement for life is likely to be quite low. Besides nutrient deficiency, early cyanobacteria also faced various types of stresses which are ‘unforeseeable’ for them. One type is the rising oxygen level they caused themselves through oxygenic photosynthesis, for which they developed various counterattack strategies in modern cyanobacteria (Raymond and Segrè 2006, Moore et al. 2017). Ultraviolet light is another early stress which has persisted until now, but some cyanobacteria evolved against it by producing sunscreens (Garcia-Pichel et al. 2019). These global stresses were also intertwined with landscape changes through geophysical processes (e.g., volcanism and climatic shifts). Here, we demonstrate the remarkable adaptive ability of cyanobacteria by showcasing their evolution of biological capability and morphology under the rising oxygen levels.

Since origin, particularly after the Great Oxidation Event (GOE) (Fig. 3), ancestral cyanobacteria quickly acquired versatile antioxidant capabilities against reactive oxygen species (ROSs), derived from O_2_ produced as a result of oxygenic photosynthesis. Different classes of superoxide dismutases (SODs) and other types of antioxidant enzymes are evolved to deal with ROSs throughout their entire history (Boden et al. 2021). Based on local availability of metal cofactors (another case of adaptation), three types of SODs—NiSOD, CuZnSOD, and MnSOD/FeSOD—have been found in four types of aquatic environments (Fig. 3A). Besides direct enzymatic removal of ROSs, cyanobacteria have also evolved other strategies, e.g., in photosynthetic reaction center (RC) which split into two types as a response to the rising oxygen (Orf et al. 2018).

**Figure 3.**
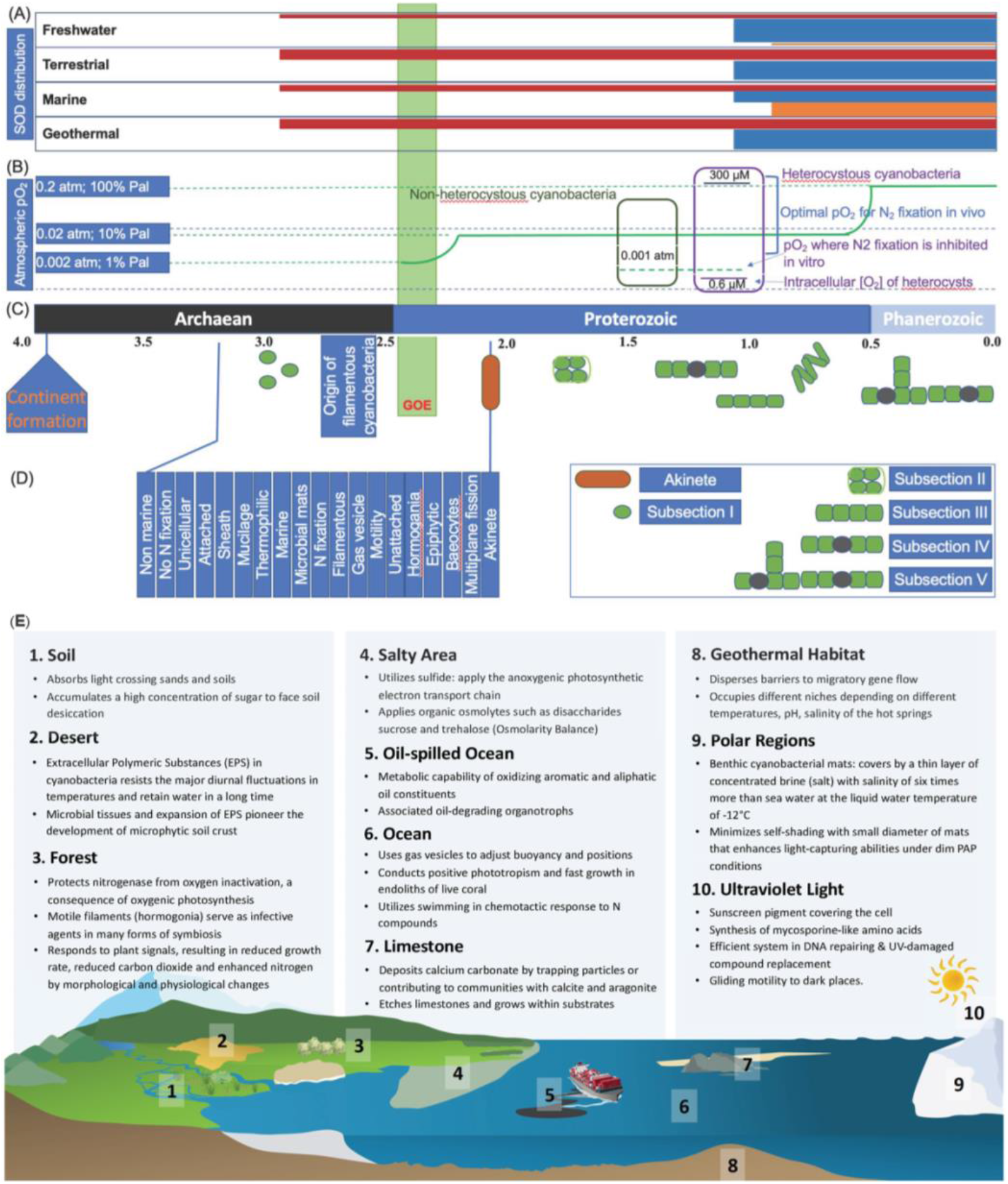
The evolutionary trajectory of cyanobacteria in the presence of oxygen and adaptive radiation. (A) The origin and distribution of SODs in different habitats. The timing of origin and habitat distribution of SOD genes among taxa is created based on Boden et al (Boden et al. 2021). Starting points of horizontal bars represent time of origin and color SOD types: CuZnSOD (blue), NiSOD (green), and Fe- and Mn-utilizing SODs (yellow). (B) The oxygen level in atmosphere and nitrogen fixation in heterocysts in cyanobacteria in the presence of elevated oxygen levels. The level of oxygen is based on (Tomitani et al. 2006). (C) and (D) The temporal morphological diversification cyanobacteria is based on Schirrmeistera et al (Schirrmeister et al. 2013). (E). Adaptive strategies of cyanobacteria in different habitats. Cyanobacterial adaptation strategies in nine environments, namely soil, desert, forest, salty area, oil-spilled ocean, open ocean, limestone, geothermal water, and polar region. Cyanobacteria also cope with the damage caused by ultraviolet light.

Another adaptation lies in the redundancy (heavy investment) of key functions. For example, in photosystem I, cyanobacteria have two electron transporters that evolved separately: copper-dependent plastocyanin (PC) and iron-dependent cytochrome c_6_ (Cytc_6_). Distinct in the primary sequence and tertiary structure of protein/amino acid, their expression is controlled by a BlaI/CopY-family transcription factor (PetR) and a BlaR-membrane protease (PetP), depending on the availability of the metal cofactors (García-Cañas et al. 2021).

With a continuous rise of oxygen level, morphological diversification was called upon to provide an extra dimension of protection. It has been confirmed that the evolution of multicellular morphotypes and the rate of morphological diversification coincide with the GOE onset (Schirrmeister et al. 2013). Heterocyst became a specialized cell form for nitrogen fixation, in which the oxygen level is reduced to pre-GOE levels (Fig. 3B). Nitrogen fixation is viewed as a leap forward in promoting marine primary production and contributed to increased O_2_ levels, coinciding with the rise of animals (Lyons et al. 2014). Similarly, cyanobacteria also obtained a specific photosynthetic structure for carbon fixation, carboxysome, in which CO_2_ is increased from 10-15µM to 40mM outside and greatly improved the efficiency of enzymatic fixation of carbon dioxide (Mangan and Brenner 2014). After the evolution of geological timescales (Fig. 3C and 3D), modern cyanobacteria are among the most diverse prokaryotic phyla, with morphotypes ranging from unicellular to multicellular filamentous forms, including those able to terminally (i.e., irreversibly) differentiate in form and function (Schirrmeister et al. 2013, Hammerschmidt et al. 2020).

Billions of years of adaptive evolution have allowed structure elaboration and functional diversification in coping with nutrient deficiency, environmental extremes and constraints (Paerl 2014). This evolutionary divergence has enabled cyanobacteria to occupy all geographic habitats in modern Earth, including terrestrial and aquatic ecosystems, ranging from deserts to tropical rain forests, soils and limestones, and from open oceans and brackish waters to freshwaters and hydrothermal vents (Fig. 3E). This broad array of biological functions underlying the adaptive radiation in cyanobacteria is recorded in their genomes (Chen et al. 2021). A large proportion of the genes in their genomes are associated with adaption to the specific habitat they occupy, but not to any other habitats. Therefore, the genomes produce a pangenome with most genes being unique and only a small set of 323 genes being common among cyanobacteria (Shi and Falkowski 2008). These core genes are mainly involved in housekeeping functions, as opposed to interacting with environment, e.g., ribosomal proteins, photosynthetic apparatus, ATP synthesis, chlorophyll biosynthesis, and the Calvin cycle. This small stable core and large variable shell of genomes suggest most of the non-overlapping functions are specialized in adaptation to distinct habitats.

Based on adaptation to rising oxygen level they produced themselves paralleling the evolution of Earth and adaptation to various habitats (see discussion above), one can conclude that most of the functions in CyanoPATH are also obtained through adaptive radiation. For example, one most prominent function is the ABC transport for phosphate PstSCAB, most cyanobacteria have two copies of operons and some top bloomers such as M. aeruginosa have three operons. Again, CyanoPATH functions are the part of entire function repertoire which sustains cyanobacteria through the harsh environment before the era of water eutrophication.

## V. The general mechanism of CyanoHABs

In light of the mechanism discussed above that CyanoHABs result from the synergistic interactions between elevated nutrients and the superior functions shaped during evolution, why it is cyanobacteria, not the eukaryotic algae, that form blooms in the same eutrophic waters? In what ways are cyanobacteria better than their co-living counterparts (e.g., Chen et al. 2017, Stockenreiter et al. 2021)? This question is so important such that the population-centric mechanism of CyanoHABs must be amended to accommodate the community-level dominance.

This issue is quite complex, considering high connectedness between populations in a food web. To reduce its complexity to a level amenable to testing without invoking any experiments, here we only consider the issue from an energy perspective. Compared to CyanoHABs as a result of maximum growth, one can inter two reasons why the co-existing algae cannot form blooms. For one, they are not able to obtain enough nutrients for growth; for the other, they have higher demands for nutrients than the current levels can provision. In the discussion below, we focus on the latter and elucidate cyanobacteria are simpler life forms than their co-living eukaryotic algae life forms and thus require less nutrients, through cell structure, size, genome size, per capita cell content, nutrient demand, and field manipulation.

### Cell Structure and size

Early deep origin and subsequent adaptive radiation (see Section IV) have shaped cyanobacteria as simplest form of life among algae. Additionally, cyanobacteria are the ancestor of eukaryotic algae, Chlorophyta, Rhodophyta, Euglenophyta, Cryptophyta, and Dinophyta commonly seen in fresh waters, e.g., in Lake Taihu (China) (Niu et al., 2011). Therefore, Cyanophyta, being a prokaryote, have the simplest cell structure among modern algae (Fig. 4A): Cyanophyta does not contain a nucleus and membrane-bound organelles such as mitochondria, Golgi apparatus, and endoplasmic reticulum. Throughout the algal evolution, diverse eukaryotic algae have gained structural complexity and acquired plastids via endosymbiotic events on their common prokaryotic ancestor, Cyanophyta (Fig. 4B). In parallel to simple structure, cyanobacteria have the smallest size among all algae gropus (Agusti et al. 1987). One example is provided with lab cultures: cyanobacterium *M. aeruginosa* and chlorophyte *Cholorella vulgaris* are 3.30 ± 0.91 μm and 5.32 ± 1.27 μm in diameter, respectively (Hu 2014).

**Figure 4.**
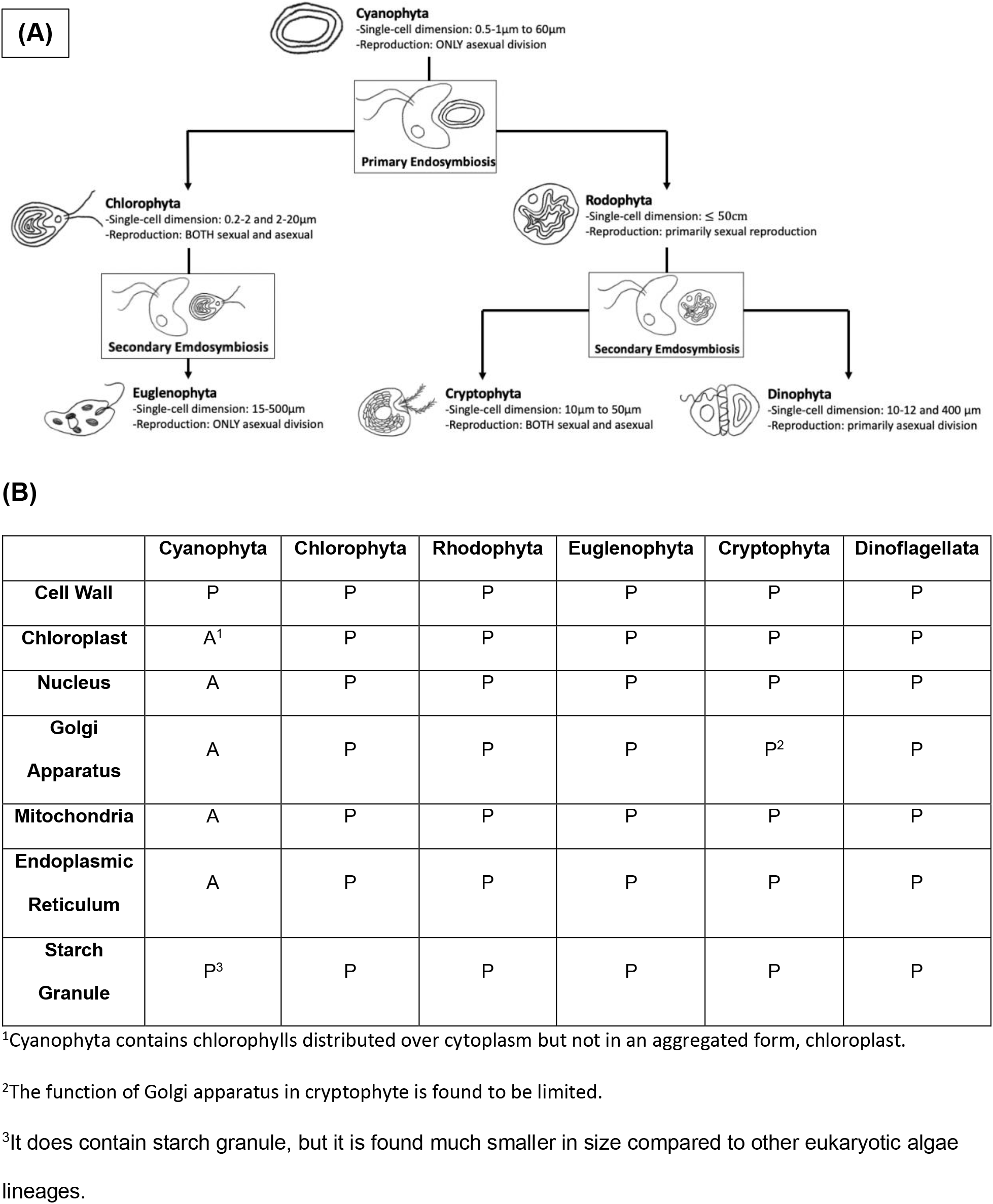
Cell structure of cyanobacteria and eukaryotic algae commonly seen in fresh waters and their evolutionary relationship. A: absent; P: present. (A) The major eukaryotic groups have risen from a common prokaryotic ancestor, Cyanophyta, via endosymbiotic events. Through the primary endosymbiosis, Chlorophyta and Rhodophyta acquired primary plastids, membrane-bound organelles usually found in eukaryotic organisms. Chlorophyta was preyed upon by a second eukaryotic cell, resulting in Euglenophyta acquiring their plastids. Similarly, Cryptophyta and Dinophyta have arisen as a result of secondary endosymbiosis on Rhodophyta. (B) Comparison of cellular components in six common major algal phyla. Cyanophyta shows the absence of nucleus, chloroplast, and mitochondria compared to other eukaryotic algae lineages. Cyanophyta, being the common prokaryotic ancestor, do not contain membrane-bound organelles.

### Genome Size

Consistent with the structural simplicity, the genome sizes of cyanobacteria are the smallest compared to those of eukaryotic algae, which are smaller by one or two orders of magnitude (Fig. 5A). Most genomes of the eukaryotic algae are about the same size of around 100 Mb base pairs, except those of Phaeophyceae and Dinophyceae being one more order of magnitude larger. Cyanobacteria usually have around 5000 genes (Kaneko et al. 2007, Frangeul et al. 2008) while eukaryotic genomes have at least twice as many (e.g., Armbrust et al. 2004, Chen et al. 2020). These differences in genome size and gene count reflect the difference in cell structure and functions.

**Figure 5.**
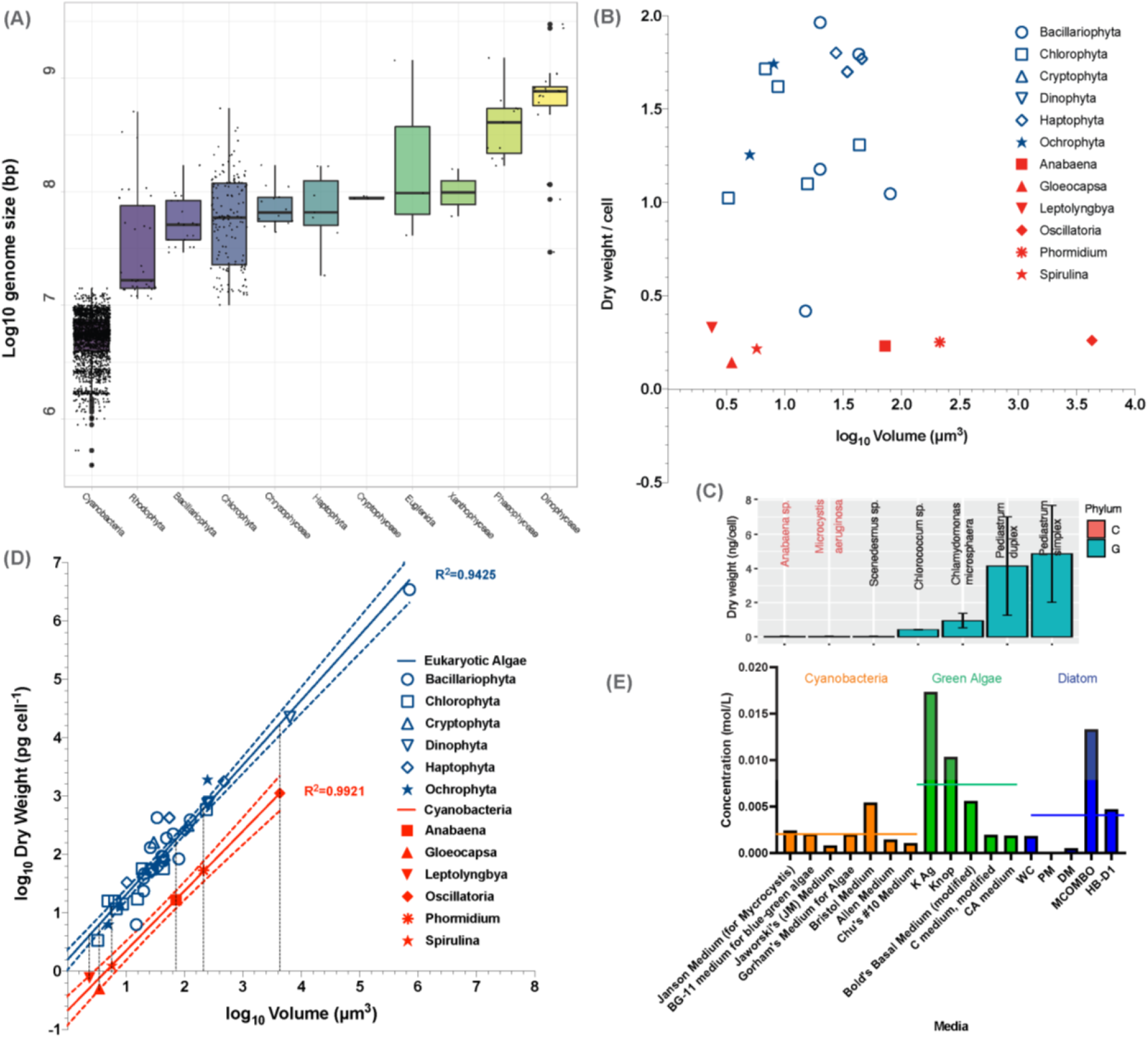
Comparison between cyanobacteria and eukaryotic algae in select key traits. (A): Distributions of log-transformed sequence length in eleven groups of algae. Summary statistics of 2240 genome assemblies were downloaded from NCBI Datasets: 1977 in cyanobacteria, 38 in Rhodophyta, 26 in Bacillariophyta, 156 in Chlorophyta, 14 in Chrysophyceae, 7 in Haptophyta, 3 in Cryptophyceae, 3 in Cryptophyceae, 3 in Euglenida, 2 in Xanthophyceae, 11 in Phaeophyceae and 20 in Dinophyceae, respectively. Each dot in the boxplot represents the log-transformed total sequence length of one assembly. (B): Distributions of dry weight per cell biovolume in Eukaryotic algae and cyanobacteria from literature (Mahlmann et al. 2008, Finkel et al. 2016). Each dot plot represents the average values of dry weight per unit biovolume and cell dry weights within a class. Blue symbols represent eukaryotic algae cells and red symbols represents cyanobacteria. (C): The per-cell dry weight of two select cyanobacteria and five green algae at mid log phase. C: cyanobacteria; G: green algae. (D): Relationship between cell volume (log_10_*V*, µm^3^), and dry weight (log_10_ *DW*, pg cell^-1^). Each dot plot represents the average values of cell volumes and cell dry weights within a class. Linear regressions were applied for eukaryotic algae (blue) and cyanobacteria (red) with 95% confidence intervals. (E): Lower nutrient requirements by cyanobacteria than eukaryotic algae in aquatic environment. Among seven selected media, the recipes of mAC, MJ, VTAC, and CHU-11 (left panel) are used for culturing green algae and the recipes of M11, BG11 and CT (right panel) are used for culturing cyanobacteria.

### Genome-Scale Metabolic Networks (GSMs)

Cyanobacteria have half the number of the genes encoded in eukaryotic genomes at the most. For some species, GSMs have been reconstructed based on the gene-protein/enzyme-reaction rule and experimental measurement of the macromolecule components (Table 1). In terms of the number of reactions and metabolites, the GSMs of cyanobacteria have, on average, 780 metabolites and 820 reactions, which are only less than half those of the eukaryotic GSMs. As the reactions are involved in catabolism, core metabolism, and anabolism (Li et al. 2018), these smaller GSMs in cyanobacteria make it clear that less nutrient input is needed than their more complex eukaryotic counterparts for life maintenance and growth. Therefore, as nutrient levels increase from the oligotrophic states during the early process of eutrophication, cyanobacteria are the first group that can be saturated for growth.

**Table 1.**
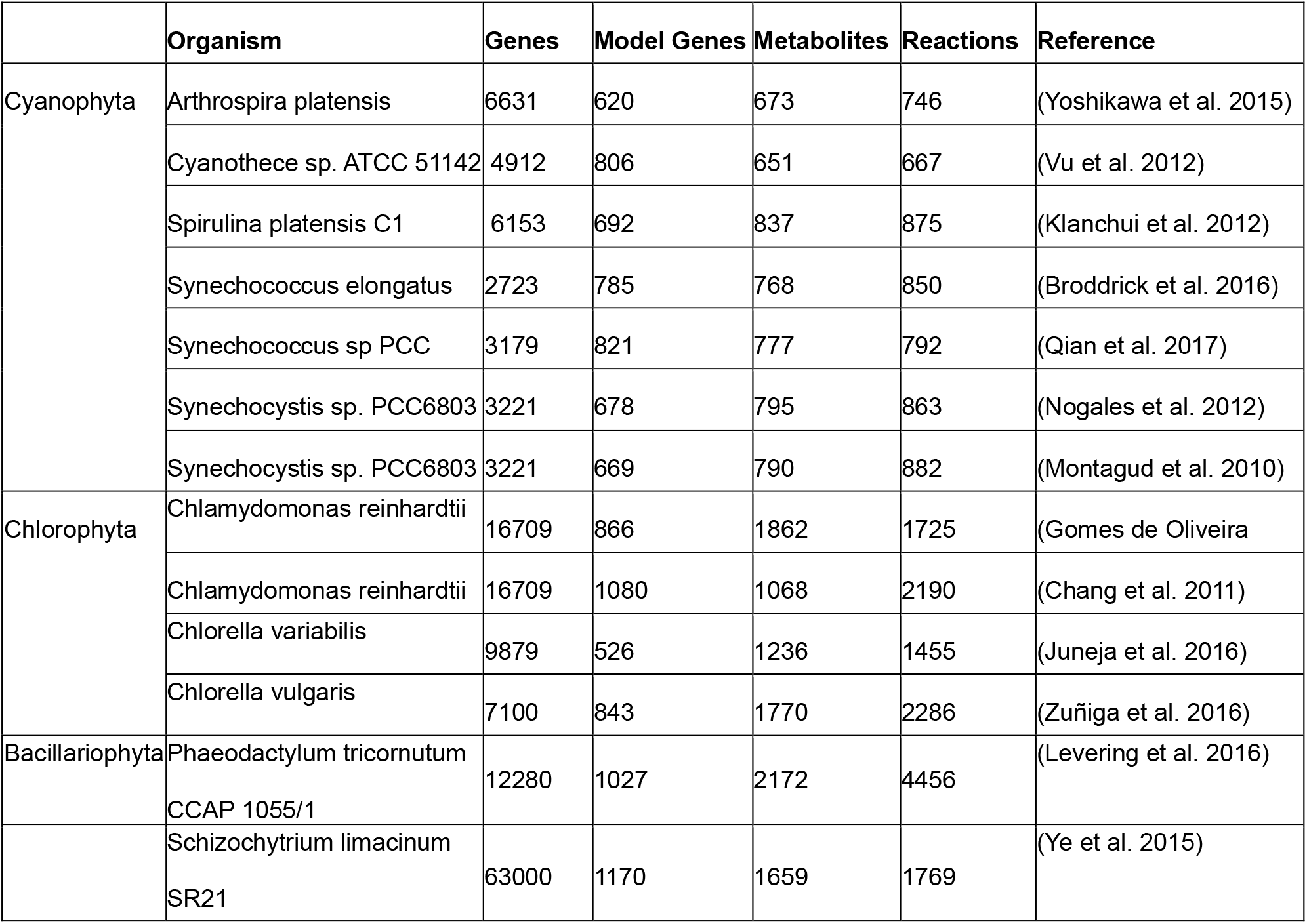
Comparison of genome-scale metabolic networks between cyanobacteria and eukaryotic algae.

Besides the size of GSMs, the metabolic rate (time^-1^) is known to often scale with cell size with a volume scaling exponent *b* = −0.25 (López-Urrutia et al. 2006, Finkel et al. 2010).

### Per capita cell content

Following the same line of life form complexity, we examined the cell content per cell between cyanobacteria and eukaryotic algae. It is clear that different species of cyanobacteria have lower cell content than those of other algae, using the data from literature (Mahlmann et al. 2008, Finkel et al. 2016) (Wilcoxon Test, *P* < 0.01; Fig. 5B). Besides the public data, we also showed that the same pattern obtained with our own per-cell cell content (as dry weight), we collected with two species of cyanobacteria and five species of green algae (Wilcoxon Test, *P* < 0.01; Fig. 5C). Interestingly, species of cyanobacteria or non-cyanobacteria show a similar trend of within-group variation in per capita cell content (Fig. 5D). Here we provide on specific example of lab cultures. Dry weight of individual *Cholorella vulgaris* cells at exponential phase are 1.70 ± 0.07×10^-11^ g/cell, and those of *M. aeruginosa* average 6.38 ± 0.05×10^-12^ g/cell, which is only about of the green alga (Hu 2014).

### Concentration of synthetic media

To further support the point that cyanobacteria are simpler life forms and require less energy and mass input, we examined the recipe of the commonly used media optimized for growing different groups of algae. As shown in Fig. 5E, the media for culturing cyanobacteria generally have lower total concentrations of ions than those for green algae or diatom, although the statistic test, Welch’s test, produced a high p value (*P* =0.1) for the selected media.

### Golden test of nutrient demand: whole-lake nutrient manipulation

To test the hypothesis that cyanobacteria have lower nutrient requirements for maximum growth than co-living eukaryotic algae, we summarized the results, which directly compared the growth of cyanobacteria with eukaryotic algae in the field, laboratory, and cosmos by supplying external nutrients (Table 2). Five case studies we discovered all support our hypothesis: low nutrients (TP, TN, DIN, urea-N, Ca, etc.) leads to dominance of cyanobacteria and high nutrients produce dominance of green algae and even brown algae.

**Table 2.**
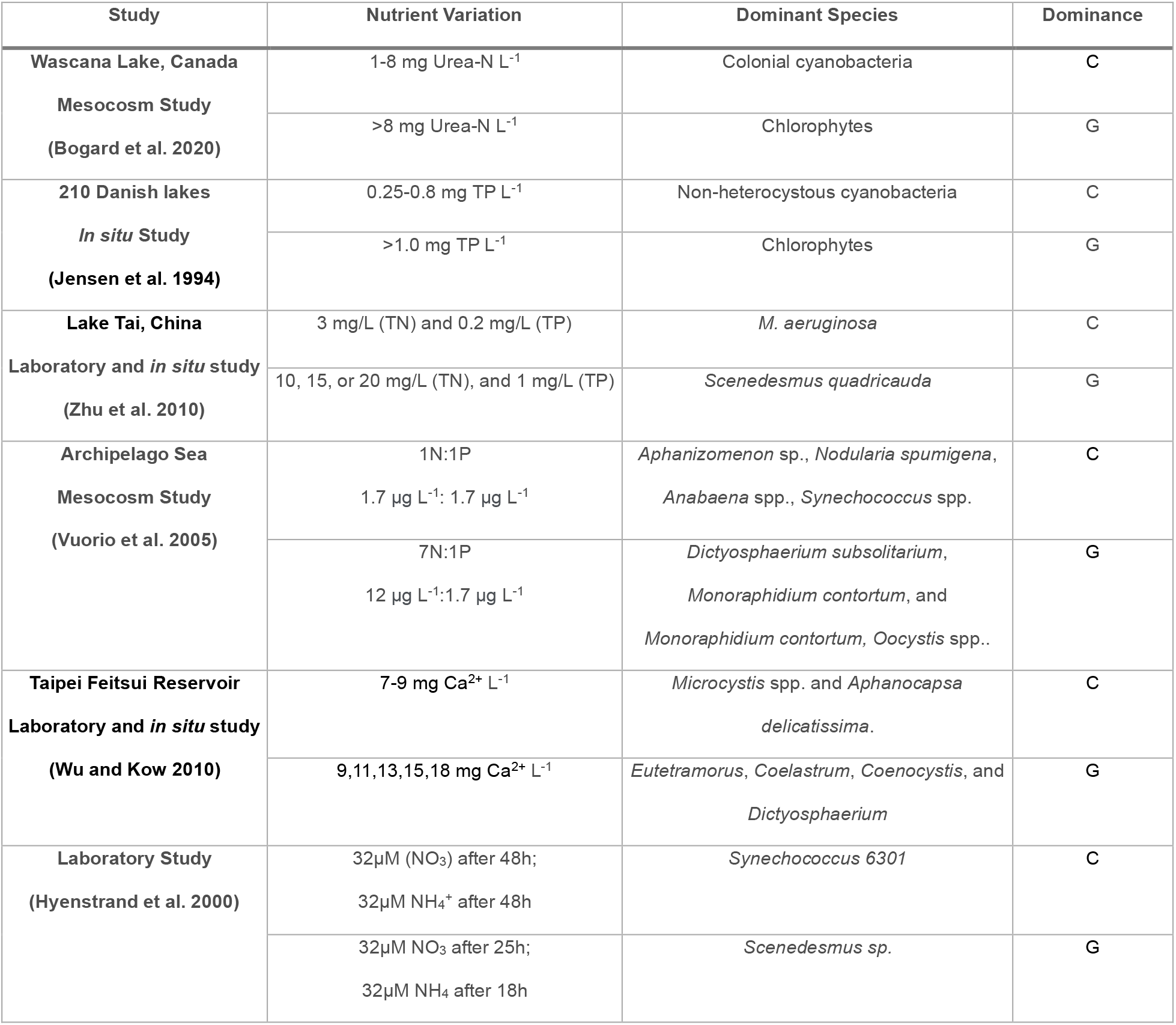
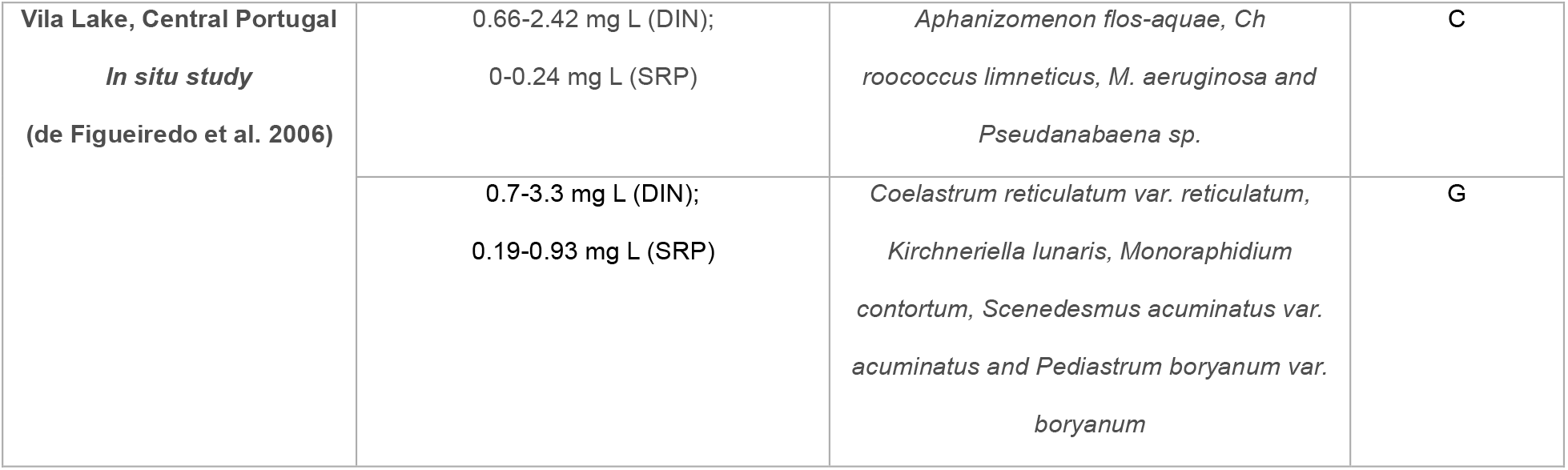
Case studies of nutrient manipulation on phytoplankton *in situ*, in the mesocosm study, and in laboratory conditions showing lower nutrient requirements of cyanobacteria than eukaryotic algae. “C” represents cyanobacteria; “G” represents green algae. DIN: dissolved inorganic nitrogen; SRP: soluble reactive phosphorus; TN: total nitrogen; and TP: total phosphorus.

In a large study of 210 Danish lakes heterocystous cyanobacteria were in dominance when the total phosphorus (TP) at low levels (<0.25 mg P·L^-1^) and non-heterocystous cyanobacteria were in dominance when TP was at intermediate levels (0.25-0.8 mg P·L-1), while chlorophytes often dominated when TP was high (>1 mg P·L^-1^) (Jensen et al. 1994),. In another whole-lake study conducted in Wascana Lake (Canada) (Bogard et al. 2020), different levels (0, 1, 3, 8, and 18 mg N L^-1^) of urea added into the surface water every seven days (day 0, day 7, and day 14) yielded compositional shifts in the phytoplankton community—after 21 days, colonial cyanobacteria dominated when N level was moderate (≤8 mg N L^-1^), and chlorophyte dominated when N level was high (>8 mg N L-1). A third support comes as a nutrient-supplement mesocosm in the Archipelago Sea (Table 2). Two ratios of N:P supplement solution, 1N (1.7 μg L^-1^):1P (1.7 μg L^-1^) and 7N (12 μg L^-1^):1P (1.7 μg L^-1^), were applied to the mesocosm at low initial concentrations of N (mean=2.7 μg L^-1^) and P (mean=2.8 μg L^-1^) every day over twenty days. At the onset, heterocystous cyanobacteria made up 70% to 80% of the total phytoplankton community. Compared to 1N:1P treatment, 7N:1P led to the most dramatic decline of the percentage of the heterocystous cyanobacteria and the chlorophyte dominance with the flourishing of *Dictyosphaerium subsolitarium*, *Monoraphidium contortum* and *Oocystis spp*. Besides N and P, the need for calcium is also considered. Lab-based calcium enrichment experiment (Wu & Kow, 2010) showed that both the cell number and the cell column of chlorophytes elevated with the increase of calcium level, in contrast to those of cyanobacteria. More examples of nutrient supplementation to shift dominance from cyanobacteria to other algae are provided in Table 2.

To sum up, cyanobacteria are simpler life forms than co-living eukaryotic algae, in cell size, cell structure, genome size, GSMs, and cell content. Simple structure and small size provide them with higher metabolic rates leading to higher abundance (Agusti et al. 1987), which have been proved by field nutrient supplementation. Now it is clear that complex eukaryotic algae would form blooms if nutrients were high enough.

## V. Synthesis and outlook

We propose a synthesis of CyanoPATH as a result of the synergistic interactions between cyanobacterial superior functions and elevated nutrients. In this evolutionary ecological perspective, we provided evidence these functions are shaped during the long course of adaptive radiation, but not toward water eutrophication. More importantly, we place CyanoHABs in the community context and show that they are simple life forms than co-living eukaryotic algae in terms of cell size, cell structure, genome size, per capita cell content, metabolic network, nutrient demand. At last, simpler cyanobacteria having low nutrient requirements are proved by various golden tests, field studies of nutrient supplement. This community-centric mechanism of CyanoHABs provides a framework to study them in a wholistic perspective, considering both the differences in energy and matter requirement of various algal cell machineries and the environment dictating their dynamics in relative to each other.

Considering the multiple factors in the mechanism, further empirical tests and theoretic explorations are needed to make a better synthesis. Two types of research should be given priority: (1) comparative studies between cyanobacteria and co-living algae in nutrient requirement and growth in lab and field settings, and their interactions with varying levels of environmental factors; and (2) the integrated systems biology assessment of the relative roles in promoting growth in cyanobacteria. The latter can be best achieved by genome-scale metabolic networks.

## Acknowledgements

This work was supported by National Natural Science Foundation of China (32171565 and 52070117) and Duke Kunshan University Summer Research Scholarships and Signature Work Research Grants (to WC, SH, QG, LQ, WL, and YY).

